# Specificity profiling of SARS-CoV-2 PLpro using proteome-derived libraries of linear peptides suggests secondary preference for basic motifs

**DOI:** 10.1101/2025.05.15.654199

**Authors:** Daniel Vogele, Klemens Fröhlich, Oğuz Bolgi, Christoph Peters, Ruth Geiss-Friedlander, Oliver Schilling

**Author notes:** Corresponding author: Prof. Dr. Oliver Schilling, Institute for Surgical Pathology, University Medical Center Freiburg, Faculty of Medicine – University of Freiburg, Breisacher Strasse 115a, D-79106 Freiburg.

## Abstract

SARS-CoV-2 papain-like protease (PLpro) is essential for viral replication and immune modulation. Here, a proteomic identification of protease cleavage sites (PICS) approach was applied using proteome-derived peptide libraries to determine the enzyme’s substrate specificity. PLpro exhibited a yet unreported preference for basic amino acids at P1, primarily arginine or lysine. This secondary specificity frequently involved cleavage between two basic residues (e.g., K|K, K|R or R|K). Experiments with in-house and commercial PLpro in GluC-generated peptide libraries from *Escherichia coli* and HEK293 proteomes confirmed this preference, though with lower overall efficiency compared to typical trypsin-like proteases. SARS-CoV-1 PLpro likewise displayed this basic-site specificity, underscoring its conservation across related coronaviruses. Site-directed mutagenesis of acidic residues near the catalytic triad to neutral variants altered cleavage preferences, indicating the involvement of these side chains in substrate binding and potential alternative binding modes. We also evaluated wild-type PLpro specificity on intact protein lysates rather than peptide libraries to assess how structure influences cleavage patterns. Notably, P1 arginine specificity persisted at the protein level, whereas lysine specificity was reduced, suggesting additional structural constraints in complex substrates. A strong presence of glycine on the prime side further suggests a bias toward unstructured regions. These findings reveal an expanded substrate recognition repertoire for SARS-CoV-2 PLpro, which may be relevant for the design of targeted inhibitors and understanding of viral protease function.

## 1. Introduction

Coronaviral replication relies on two cysteine proteases, the main protease (Mpro) and the papain-like protease (PLpro). For Mpro already there exists the clinically approved inhibitor nirmatrelvir [1], while therapeutic PLpro inhibition is not being pursued. This is despite PLpro performing a dual task: it cleaves three junctions in the viral polyprotein (nsp1|2, nsp2|3, nsp3|4) and antagonises innate immunity by removing ubiquitin and ISG15 from host signaling proteins [2–4]. Both activities hinge on a catalytic triad (Cys111-His272-Asp286 in SARS-CoV-2 numbering) and a minimalist tetrapeptide consensus, LXGG↓X, first characterised for SARS-CoV-1 and later confirmed for SARS-CoV-2 [5–7]. Because LXGG mimics the C-terminal tail of ubiquitin and ISG15, most biochemical studies have focused on viral junctions or fluorogenic peptides that present the di-glycine motif.

Despite this perceived stringency, scattered evidence suggests that PLpro may recognise additional sequence classes. SARS-CoV-2 PLpro can hydrolyse Lys48-linked poly-ubiquitin chains whose isopeptide junction displays RLRGG [4], and N-terminomic analyses of infected cells have reported host cleavage sites that deviate from LXGG [8]. However, those studies relied either on overexpression systems or trypsin-generated peptide libraries, which lack internal lysine and arginine and thus cannot expose preferences for basic residues at P1 or P1′. Moreover, the structural basis for any extended motif and its conservation between SARS-CoV-2 and the closely related SARS-CoV-1 PLpro remain unexplored. Consequently, the full breadth of PLpro specificity and its mechanistic basis remain unsolved. Defining these features is essential, because putative PLpro inhibitors should block both viral polyprotein processing and cleavage of host substrates that modulate immunity.

Proteomic Identification of Cleavage Sites (PICS) provides an unbiased means to profile protease specificity from thousands of sequences simultaneously [9]. Crucially, when peptide libraries are generated with GluC rather than trypsin, internal basic residues are retained, allowing full assessment of prime- and non-prime-side requirements. Quantitative terminomic methods such as TAILS or *in silico* terminomics complement PICS by mapping cleavage events in folded proteins and thereby incorporating structural context [10,11]. Combining these approaches with isobaric labelling yields a unified platform to interrogate specificity from peptide mixtures to native lysates.

In the present study, we use GluC-based PICS, tandem-mass-tag quantification, and a terminomics workflow on intact cell lysates to map the substrate preferences of SARS-CoV-2 PLpro. We benchmark its motif against that of SARS-CoV-1 to assess evolutionary conservation; titrate enzyme-to-substrate ratios and compare turnover with trypsin to place the activity in biochemical context; and employ site-directed mutagenesis of three acidic residues (Asp108, Glu161, Asp164) that border the substrate-binding cleft to probe their role in extended recognition. By integrating peptide-level and protein-level datasets under a single analytical framework, our work provides an expanded sequence template for PLpro, delineates structural features that govern basic-site recognition, and supplies information critical for the rational design of inhibitors capable of blocking both viral and host-directed cleavage events.

These insights address a key gap in coronavirus biology and may inform therapeutic strategies based on viral protease specificity.

## 2. Materials and methods

### 2.1. Generation of in-house wild-type and site-mutant PLpro

The cloning and purification strategies for both wild-type (WT) PLpro and its site-directed mutants were adapted from Steuten et al. (2020) [12]. In brief, PLpro was fused to an N-terminal 6×His-SUMO1 tag and expressed in *Escherichia coli* BL21 (DE3). The resulting protein was bound to Ni-NTA resin (Qiagen), and the 6×His-SUMO tag was subsequently cleaved using GST-SENP, yielding untagged PLpro with its native N-terminus. The protease was concentrated using an Amicon 5 kDa centrifugal filter and further purified by gel-filtration. To generate PLpro variants D107N, E160Q, E163Q, and the active-site mutant C111A, point mutations were introduced by site-directed mutagenesis. Each mutation replaced the corresponding residue with asparagine (Asp→Asn), glutamine (Glu→Gln), or alanine (Cys→Ala) at their respective position referring to the PLpro amino acid sequence.

### 2.2. PICS workflow

#### 2.2.1. Peptide library preparation

HEK293 or *E. coli* cell pellet were resuspended in library digestion buffer (0.1% Rapigest, 100 mM HEPES) and cell lysis was performed exclusively by ultrasonication with a Bioruptor for 20 cycles (40 s on, 30 s off) at a high intensity setting. After a 10-minute centrifugation at 15,000×g, the supernatant was collected and the protein concentration determined. Approximately 1 mg of protein underwent reductive alkylation using 5 mM TCEP and 20 mM IAA. GluC (Promega Corporation) digestion proceeded in two steps: first, 1/50th of the total protein amount was added as GluC, followed by a 2-hour incubation at 42°C. Then, another 1/50th of GluC was added, and the mixture was incubated overnight at 37°C. The resulting GluC-specific peptide library was desalted using Sep-Pak C18 cc Vac Cartridges (Waters™) according to manufacturer’s instruction, then vacuum dried.

#### 2.2.2. Protease digestion of libraries

Dried peptides were resuspended in digestion buffer containing either 100 mM HEPES (pH 8) or sodium phosphate buffer (pH 6), as well as 150 mM NaCl and 5 mM DTT. Key parameters of each PICS experiment, such as digestion buffer, proteases, controls and number of replicates are listed in **Supplemental Table S1**. Peptide concentration was determined and the peptide library was divided depending on the subsequent experimental procedures. For the TMT-based titration/trypsin and site-mutant experiments, the peptide library was devided into 16 samples of 25 µg peptide each. For all other heavy/light NHS-acetate-labeling experiments, the library was divided into six samples of 30 µg. Proteases of interest and negative controls were added in their respective ratios and replicates, and in case of PLpro, it was preincubated for 30 min at 37°C in 5 mM DTT (**Supp. Table S1**). Following an overnight incubation at 37°C, protease activity was halted by boiling for 10 min at 95 °C.

#### 2.2.3. NHS-acetate labeling

Buffer was adjusted to pH 8 and 100 mM HEPES for labeling using light and heavy N-hydroxy succinimide acetate (NHS-acetate). Heavy NHS-acetate was synthesized using isotopically labeled acetic acid (^13^C for one carbon and deuterium replacing three hydrogens of the methyl group), resulting in an overall mass shift of 4.01 Da. Three milligrams of either light or heavy NHS-acetate was added to the 30 µg of treated peptide library. Heavy NHS-acetate was used for the active PLpro-treated samples, while light NHS-acetate was used for the controls, with a label switch in each last replicate. Samples were incubated for 2 h at room temperature and quenched by adding hydroxylamine to a final concentration of 2%. Heavy- and light-labeled samples were combined per replicate and desalted using Hypersep™ SpinTip C-18 (Thermo Scientific) according to manufacturer’s instructions using a modified wash buffer containing 10% acetonitrile and a stepwise elution for HEK293 proteome-based experiments. Eluates collected at 10%, 14%, 18%, 22%, 26%, 30%, 40%, and 65% acetonitrile were combined pairwise (10%+26%, 14%+30%, 18%+40%, 22%+65%) to generate four fractions per replicate. All eluates were vacuum dried and submitted for mass spectrometric measurement.

#### 2.2.4. TMT labeling and fractionation

For TMT labeling, the buffers of peptide library were adjusted to pH 8 and 150 mM HEPES. Then, 200 µg of TMT reagent was added to the treated peptide libraries, aiming for a 1:8 peptide-to-reagent ratio (**Supp. Table S2**). Samples were incubated at room temperature for 3 h, followed by overnight incubation at 37°C (500 rpm). Reaction was then quenched by adding hydroxylamine to a final concentration of 2%, incubated for 30 minutes, combined into one TMT set, and vacuum dried. Samples were resuspended in high-pH buffer A (10 mM ammonium formate, pH 10) and ∼160 µg of peptide was submitted to high-pH fractionation on an Agilent 1100 HPLC system with a XBRIDGE® Peptide BEH C18 column (3.5 μm, 130 Å, 1×150 mm) at a flowrate of 40 µL/min. A 60-minute gradient from 10 mM ammonium formate (pH 10.0) to 70% acetonitrile (10 mM ammonium formate, pH 10). Total of 48 fractions were collected and concatenated into 12 fractions before MS measurement.

### 2.3. Lysate digestion and *in silico* terminomics

HEK293 cells were resuspended in 20 mM HEPES and lysed using both a syringe-and-needle method and ultrasonication (Bioruptor, 20 cycles, 40 s on, 30 s off, high intensity). After centrifugation at 15,000×g for 10 minutes, the supernatant was collected and protein concentration was measured. Eight replicates of 25 µg protein were prepared, with buffer adjsutment to 100 mM HEPES and 5 mM DTT. Active commercial PLpro (R&D systems E-611) was added to four of the replicates and all samples were incubated at 37°C overnight. Protease was inactivated by boiling at 95°C for 10 minutes. Unless stated otherwise, subsequent steps were performed as described in the previous sections. TMT reagents were added, incubated and quenched; samples were pooled, reductively alkylated and digested with GluC (Promega Corporation) at a 1:10 peptide-to-enzyme ratio. The final mixtures were desalted with Hypersep™ SpinTip C-18 (Thermo Scientific, vaccum dried, and subjected to high-pH fractionation. Fractions 8-48 were concatenated into 10 fractions before MS measurement.

### 2.4. Mass spectrometry

For MS measurement, vacuum dried peptides were solubilized in 0.1% (v/v) formic acid, sonicated for 5 min and centrifuged at 20,000g for 10 minutes before transferring the supernatant to a measurement tube. Eight hundred nanograms of each sample, together with 200 fmol of indexed retention time (iRT) peptides, were analyzed as previously described [13], using a Q-Exactive plus (Thermo Fisher Scientific) coupled to a nanoflow liquid chromatography (LC) system Easy-nLC 1000 (Thermo Fisher Scientific). The mass spectrometer was operated in data-dependent acquisition mode with a Top10 method, where each MS scan was followed by up to 10 MS/MS scans. The mass range 300-2000 m/z was recorded at a MS1 resolution of 70,000, with an automatic gain control (AGC) target of 3e6 and a maximum injection time of 50 ms. MS2 scans were acquired at a resolution of 35,000, AGC of 1e5, and a 100 ms maximum injection time, using stepped normalized collision energy (NCE) of 32.

### 2.5. Data analysis

Raw data were analyzed using the FragPipe pipeline (v22.0) [14–16] with a *E*.*coli* or human proteome database downloaded from Uniprot on March 10, 2025 (4,519 *E*.*coli* and 82,553 human entries). Decoys and contaminants were added via FragPipe’s database configuration. Searches used a 10 ppm precursor mass tolerance and 20 ppm fragment mass tolerance, with a semi-specific GluC cleavage specificity (up to three missed cleavages). Carbamidomethyl at cysteines was set as fixed modification for all analysis. For PICS TMT data, a TMT delta mass of 229.16293 Da at lysines and N-termini was set as fixed. For *in silico* terminomics TMT data, N-terminal acetylation and N-terminal TMT were set as variable modifications instead. Heavy/light NHS-acetate data used mass deltas of 46.03263 and 42.0106 as variable modification at lysines and N-termini.

FragPipe output files were further processed and analyzed in R (v4.1.0) within Rstudio. For NHS-acetate data, files were filtered, contaminants removed and semi-specific peptides with log_2_ fold-changes > 4 were identified. This set of peptides was normalized to typical amino acid abundance and cleavage motifs were displayed as heatmaps.

PSM files of TMT-based data were filtered by purity, probability, and coefficient-of-variation criteria, then processed with a modified TermineR method [11] to extract endogenous proteolytic processing information. iRT intensities were used to validate TMT channel assignments. Principal component analysis (PCA) and partial least squares discriminant analysis (PLS-DA) were performed using the mixOmics package (v6.26.0). Peptides were annotated using the TermineR package (v1.0.0) or Fragterminomics package (v0.2.2) and differential abundance analysis of semi-specific peptides was conducted using limma (v3.58.1). Their cleavage patterns were normalized to typical amino acid frequencies and visualized with pheatmap (v1.0.1).

## 3. Results and discussion

### 3.1. E. coli proteome-based PICS profiles

PICS specificity profiles were acquired using GluC peptide libraries digested by both in-house and commercially available PLpro in different pH buffers. The present study focuses on GluC specific peptide libraries because typically used tryptic PICS libraries lack internal lysine and arginine and therefore do not allow to profile specificity of basic residues in the P1 and P1’ position. Initially, *E. coli* proteome-based peptide libraries were used to profile in-house and commercial PLpro under pH 6.0 and pH 8.0 conditions. For in-house PLpro, the active-site C111S variant was used as a control, while commercial PLpro reactions were paired with a no-enzyme control. Semi-specific peptides with a log_2_ fold-change > 4 (equivalent to a 16-fold enrichment as compared to control) were used to assess positional occurrence of amino acids, resulting in 85 to 287 peptides per condition (**Fig. 1**).

**Figure 1:**
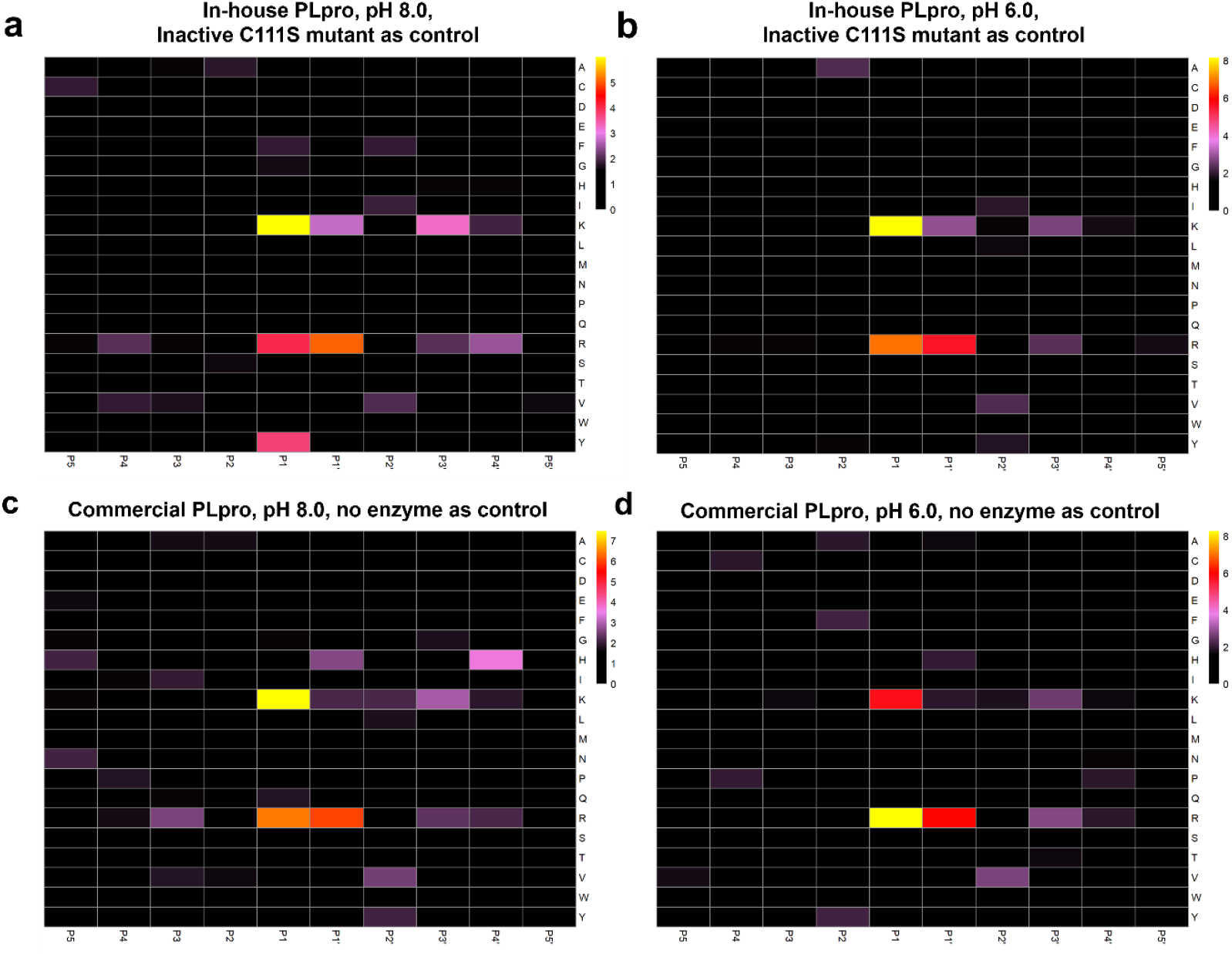
PICS specificity profiles for PLpro using *E*.*coli* proteome derived peptide libraries. (**a**) In-house PLpro incubated at pH 8.0 using C111A active site mutant as negative control (cleavage sequences n = 140). (**b**) In-house PLpro incubated at pH 6.0 using C111A active site mutant as negative control (cleavage sequences n = 287). (**c**) Commercial PLpro incubated at pH 8.0 using no PLpro addition as negative control (cleavage sequences n = 85). (**d**) Commercial PLpro incubated at pH 6.0 using no PLpro addition as negative control (cleavage sequences n = 177). PICS specificity profiles for P5-P5’ were determined as described in materials and methods. Positional occurrence of amino acids are shown as enrichment over expected frequency with natural amino acid abundance derived from Uniprot *E. coli* reference proteome UP000000625. Only enrichments >1.5-fold are displayed.

In order to distinguish genuine subsite specificity from random fluctuations, normalized amino acid frequencies were visualized only when enriched by at least 1.5-fold relative to natural abundance (lysine: natural abundance 4.41 % → cutoff 6.62 %; arginine: natural abundance 5.52 % → cutoff 8.28 %). Overall, these data revealed notable enrichment for basic amino acids (lysine and arginine) in the P1 and P1′ sites across all conditions. As an additional control, we also probed the specificity of commercially available PLpro enzyme using no enzyme addition as control. A very similar specificity profile was found; including the predominant preference for basic residues in the P1 and P1′ sites. A higher number of peptides surpassed the significance threshold at pH 6.0 than at pH 8.0, indicating elevated PLpro activity at lower pH (in-house: 287 vs. 140; commercial: 177 vs. 85). These findings guided subsequent assays using human cell-derived libraries to confirm and refine the observed substrate preferences of PLpro.

### 3.2 HEK293 proteome-based PICS profiles and comparison to SARS-CoV-1

To extend the findings observed with *E. coli* proteome-derived peptide libraries, a similar PICS approach was applied to HEK293 cell-derived proteome libraries, generated by GluC digestion. The larger sequence space available in HEK293 libraries yielded a higher number of unique potential cleavage sites (3328, pH 6.0, commercial PLpro, no enzyme addition as control, providing an expanded view of PLpro’s substrate preferences (**Fig. 2a**). Overall, the P1 enrichment for basic amino acids (lysine and arginine) persisted, confirming the preference observed in the *E. coli* experiments. Compared to the *E. coli* libraries, the relative frequency of peptides carrying a lysine at the P1′ position was lower. Arginine was the most frequent residue at the P1 site, followed by a pronounced enrichment for arginine at the P1′ position and a secondary preference for lysine in P1 (**Fig. 2a**). A minor preference for alanine was also noted at P1′, although it remains close to background activity. These observations suggest that the fundamental specificity profile remains stable, even when moving from bacterial to human proteomes.

**Figure 2:**
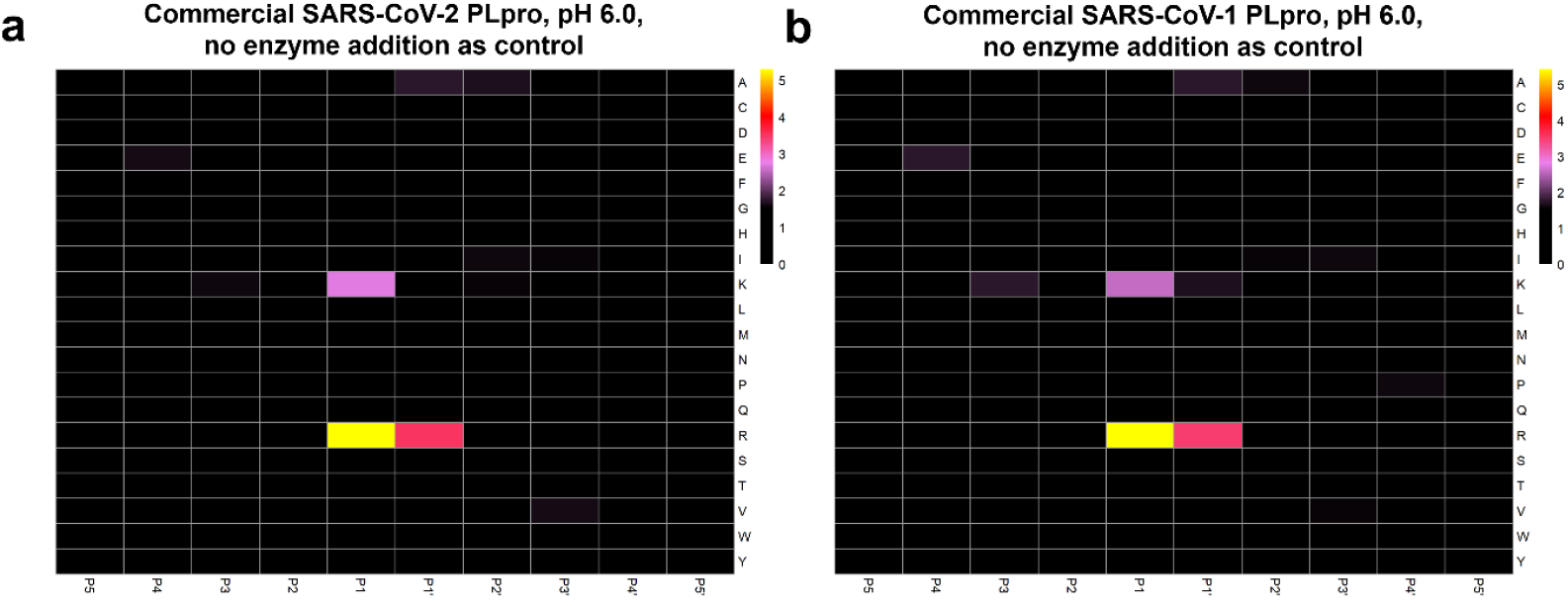
PICS specificity profiles for PLpro using HEK293 proteome derived peptide libraries. (**a**) commercial SARS-CoV-2 PLpro incubated at pH 6.0 using no PLpro addition as negative control (cleavage events n = 3328). (**b**) commercial SARS-CoV PLpro incubated at pH 6.0 using no PLpro addition as negative control (cleavage events n = 3097). PICS specificity profiles for P5-P5’ were determined as described in materials and methods. Positional occurrence of amino acids are shown as enrichment over expected frequency with natural amino acid abundance derived from Uniprot *homo sapiens* reference proteome UP000005640. Only enrichments >1.5-fold are displayed.

In a parallel experiment, SARS-CoV-1 PLpro (sourced commercially) was profiled under the same conditions and displayed nearly identical site-specific frequencies (**Fig. 2b**). This similarity is consistent with an alignment of the PLpro sequences from SARS-CoV-1 and -2 (Uniprot IDs P0C6×7 and P0DTC1, respectively) highlighting a very high degree of sequence conservation (**Suppl. Fig. 1**).

Taken together, these data reaffirm PLpro’s ability to cleave substrates at basic sites, with arginine featuring more prominently than lysine in the HEK293 libraries.

### 3.3 PLpro titration

Next, we sought to investigate the cleavage efficiency of PLpro towards a proteome-derived, GluC-generated peptide library. To this end, we assayed different PLpro:library ratios (wt/wt; 1:1000, 1:100; 1:10) as well as probing the protease trypsin for comparison (trypsin:library ratio of 1:100 (pH 8.0, over-night digestion, no enzyme addition as control). We employed multiplexed, isobaric labelling (tandem mass tags, TMT) (**Fig. 3**). For trypsin, 2015 cleavage sequences were identified. The PLpro titration yielded no cleavage sequences at the ratio of 1:1000, 244 cleavage sequences at the ratio of 1:100, and 1433 cleavage sequences at the ratio of 1:10 (**Fig. 3d**). These numbers indicate that the cleavage efficiency of PLpro towards the proteome-derived peptide library is substantial but does not reach the level of the trypsin. The resulting PLpro specificity profiles revealed arginine as the dominant residue at both P1 and P1′ positions, and a lesser but evident preference for lysine in P1. These observations are consistent with the preference for basic residues seen in the previous experiments, although the dataset included a range of background cleavage events (**Fig. 3a-b**). Trypsin displayed the classic preference for arginine and lysine at P1 (**Fig. 3c**), with the absence of an arginine preference in P1’ distinguishing tryptic from PLpro-mediated cleavage patterns. Overall, these titration data confirm that PLpro cleaves at basic residues in P1 and P1’ in a concentration-dependent manner and with a modest activity compared to the canonical serine protease like trypsin.

**Figure 3:**
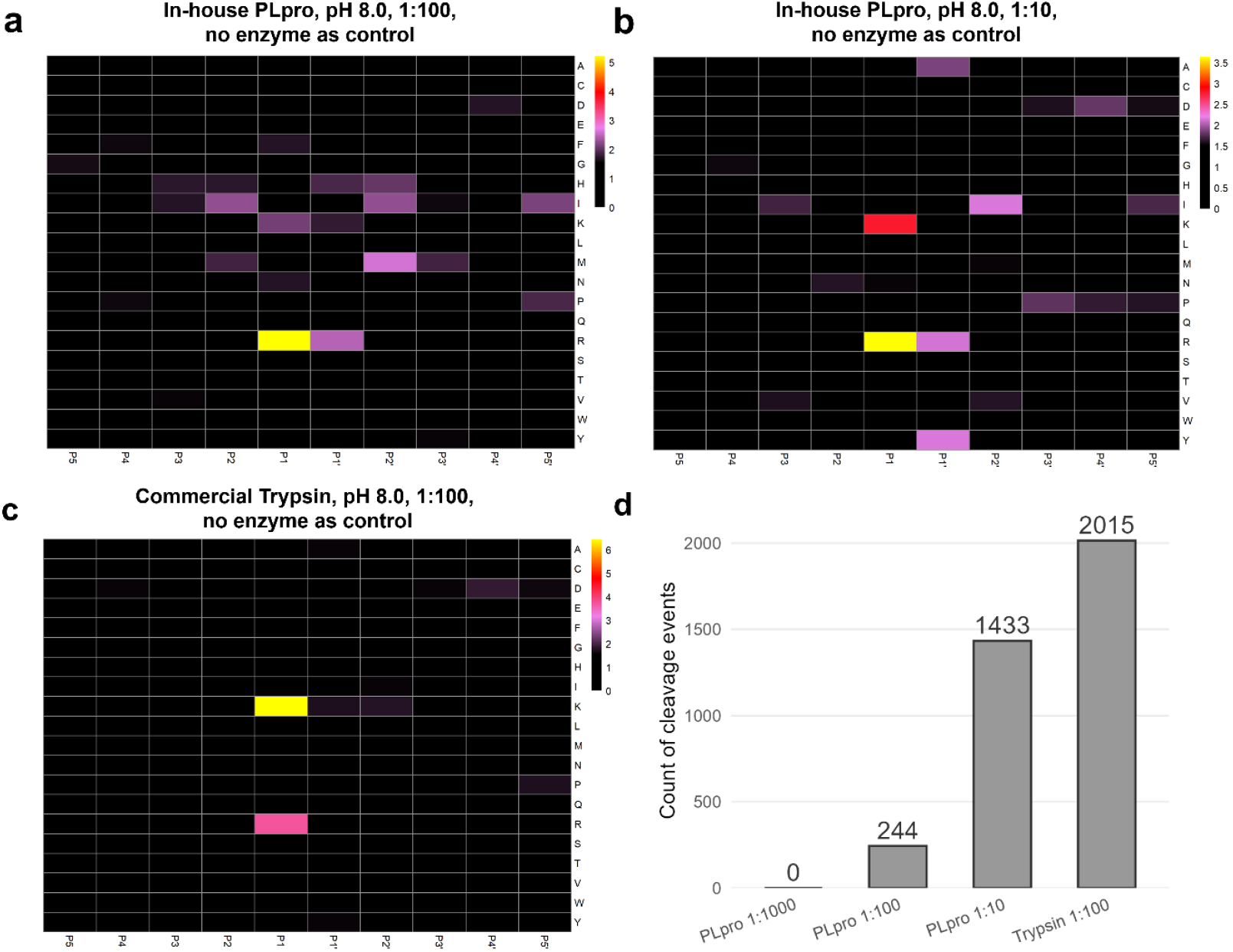
PICS specificity profiles for PLpro titration and trypsin using HEK293 proteome derived peptide libraries. (**a**) In-house PLpro incubated at pH 8.0 in a 1:100 peptide to PLpro ratio (cleavage events n = 244). (**b**) In-house PLpro incubated at pH 8.0 in a 1:10 peptide to PLpro ratio (cleavage events n = 1433). (**c**) Trypsin incubated at pH 8.0 in a 1:100 peptide to PLpro ratio (cleavage events n = 2015). (**d**) Summary bar plot showing the total number of cleavage events detected under the three conditions in panels a–c. PICS specificity profiles for P5-P5’ were determined as described in materials and methods. Positional occurrence of amino acids are shown as enrichment over expected frequency with natural amino acid abundance derived from Uniprot *homo sapiens* reference proteome UP000005640. Only enrichments >1.5-fold are displayed.

### 3.4. Acidic site-mutants and sub-site cooperativity

Having established PLpro’s preference for basic substrates, we asked whether acidic residues adjacent to the catalytic triad could stabilise arginine- or lysine-containing peptides. The crystal structure of the C111S variant (**Fig. 4a**) shows three such residues (Asp-108, Glu-161 and Asp-164) clustered near the active site. By site-directed mutagenesis, we generated three Plpro variants, each containing the carboxamide variant of the acidic residue (D107N, E160Q or E163Q, respectively). The resulting PLpro variants were examined alongside wild-type (WT) PLpro with a HEK293 GluC peptide library at pH 8.0 (enzyme:library 1:5 wt/wt), using a multiplexed TMT-based workflow for direct comparability and no enzyme addition as control (**Fig. 4**).

**Figure 4:**
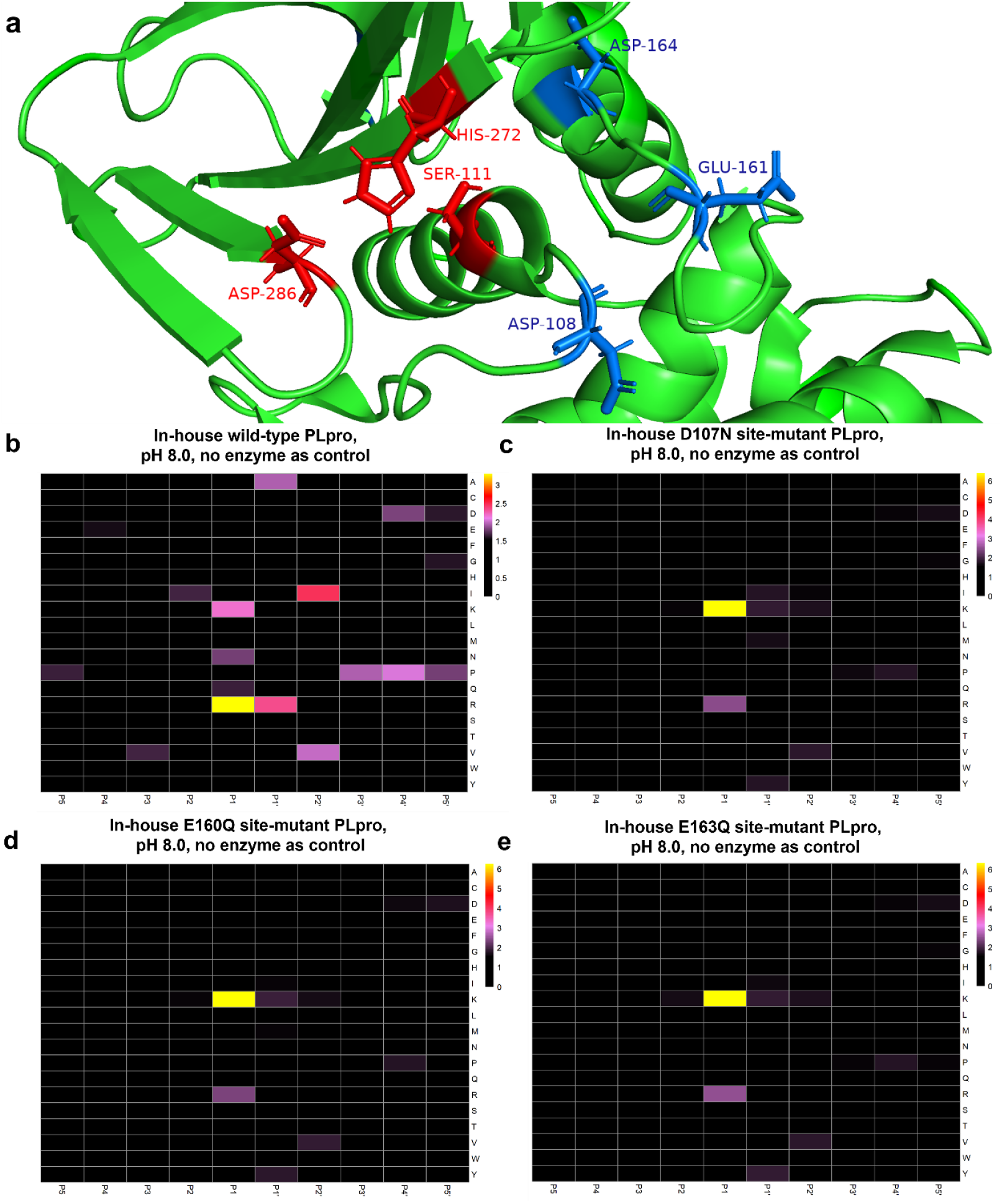
Crystal Structure of the SARS-CoV-2 PLpro C111S Mutant (PDB 7CJD) and PICS specificity profiles for wild-type PLpro and site mutants using HEK293 proteome derived peptide libraries. (**a**) This three-dimensional view of the C111S mutant (green) highlights the catalytic triad residues (Asp-286, His-272, and Ser-111) in red. Ser-111 corresponds to the mutated position of Cys-111 in the wild-type enzyme. Nearby acidic residues (Asp-108, Glu-161, and Asp-164) are shown in blue, illustrating their proximity to the active site and representing the positions selected for the individual D107N, E160Q, and E163Q mutations. (**b**) in-house wild-type PLpro incubated at pH 8.0 (cleavage events n = 1096). (**c**) in-house D107N site mutant PLpro incubated at pH 8.0 (cleavage events n = 1661). (**d**) in-house E160Q site mutant PLpro incubated at pH 8.0 (cleavage events n = 1652). (**e**) in-house E163Q site mutant PLpro incubated at pH 8.0 (cleavage events n = 1679). PICS specificity profiles for P5-P5’ were determined as described in materials and methods. Positional occurrence of amino acids are shown as enrichment over expected frequency with natural amino acid abundance derived from Uniprot *homo sapiens* reference proteome UP000005640. Only enrichments >1.5-fold are displayed.

The WT enzyme yielded 1,096 peptidic cleavage products, displaying the same di-basic motif reported earlier: arginine was strongly favoured at P1 and P1′, with lysine appearing as a secondary preference at P1 (**Fig. 4b**). Each mutant, by contrast, yielded between 1,652 and 1,679 cleavage events, indicating that proteolytic activity towards the peptide library was retained (**Fig. 4c–e**). All three variants displayed a consistent specificity shift: enrichment of arginine on the prime side disappeared entirely, and the dominant residue at P1 switched from arginine to lysine. Intersection analysis (**Suppl. Fig. 2**) reveals that only 236 cleavage sequences are shared between WT and site-mutants. 788 sequences are unique to the WT and 1218 are shared exclusively amongst the three mutants, indicating that WT and mutant enzymes cleave rather dinstinct repertoires of peptides. This surprising result remains difficult to interpret. It appears that the acidic residues under investigation likely contribute to the prime site specificity. The similar effect of all three variants may point to a (now disrupted) intramolecular network forming the structural basis for accommodating basic residues in P1’. The retained - or slightly increased - overall activity of the PLpro variants may suggest improved “accessibility” of the active site, perhaps stemming from reduced structural hindrance or selectivity in the prime site subsites. At the same time, the structural basis for non-prime site specificity remains elusive.

We further assessed putative sub-site cooperativity of the WT enzyme and the carboxamide variants for P1K, P1R, P1′K, and P1′R (**Fig. 5** for WT and D107N; full comparison of all mutants in **Suppl. Fig. 3**). The plots indicates which residues are over- or under-represented relative to their global abundance when a basic residue occupies P1 or P1′. Major subsite cooperativity (e.g. difference by > percentage points) was absent. For WT and variants, P1K and P1R were associated with a mild increase of P1’K, indicating that the preference for basic residues in P’K is not fully abolished in the carboxamide variants.

**Figure 5:**
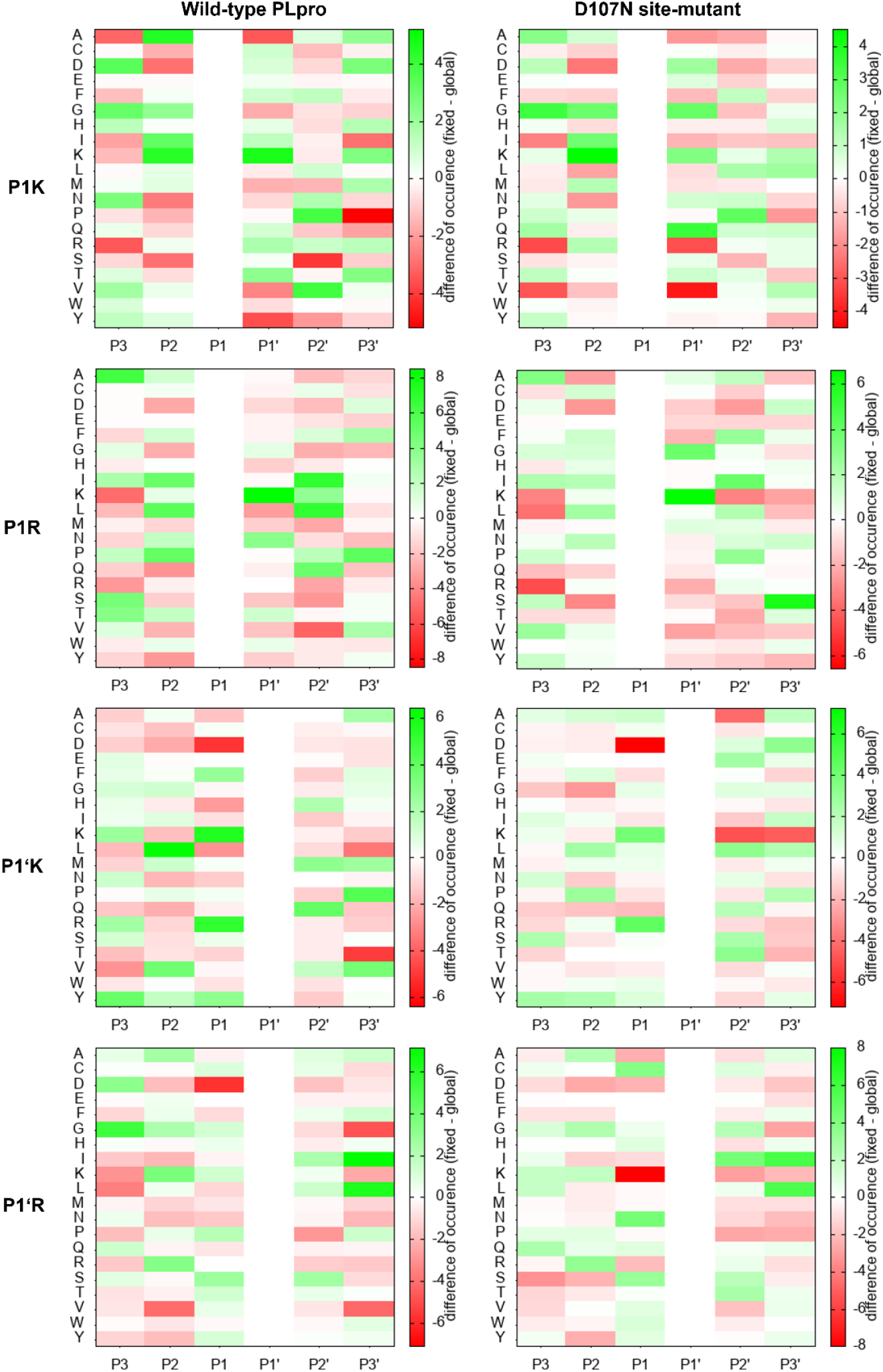
Probing for putative sub-site cooperativity for SARS-CoV-2 PLpro (WT) and representative D107N site-mutant cleavage sequences with basic residues in P1 and P1’. Each subpanel depicts subsite cooperativity analysis for wild-type SARS-CoV-2 PLpro and the D107N site-mutant in cleavage sequences containing basic residues (K or R) in P1 and P1′. Rows represent amino acids, columns indicate positions (P3 to P1, P1′ to P3′), and the color scale indicates differences from a global reference frequency (red: depletion, green: enrichment).

### 3.5. Protein-level PLpro cleavage in native HEK293 cell lysates

We sought to complement the specificity data based on peptide libraries with PLpro cleavage of native proteins as these represent longer putative substrates with a folded structure. We harvested the proteome of cultured HEK293 cells by mechanical lysis to preserve protein structure. Incubation with PLpro (pH 8.0, overnight, 1:5, no enzyme addition as control) and subsequent N-terminomics [11] using GluC digestion yielded only 59 cleavage sequences. The preference for basic residues in P1 or P1’ remained evident (**Fig. 6**). The strongly decreased number of cleavage sequences as compared to peptide-level cleavage assays suggests a PLpro preference for short and/or unfolded substrates. On the prime side, a preference for glycine residues emerges in P3′–P5′. This may be related to a preference for structural flexibility in the vicinity to the actual cleavage site.

**Figure 6:**
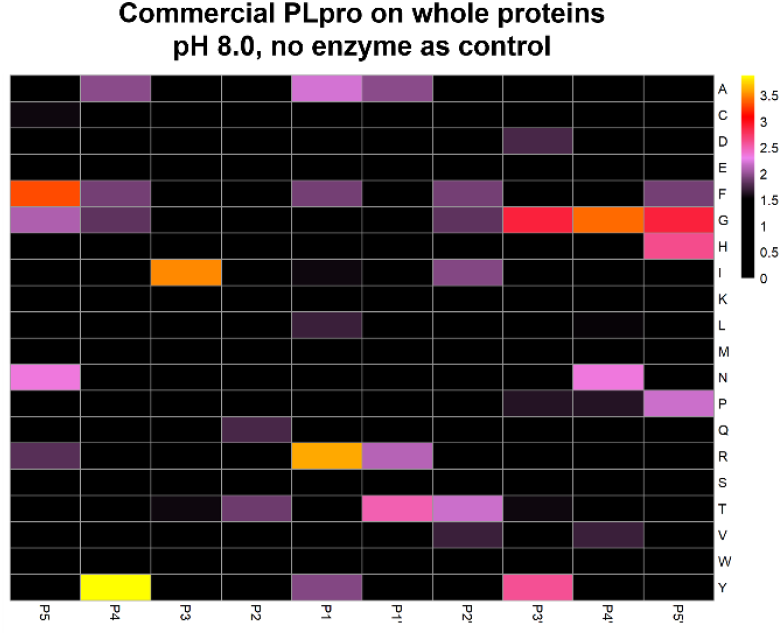
*In-silico* terminomics-derived specificity profiles for PLpro in native HEK293 cell lysate. Commercial SARS-CoV-2 PLpro incubated at pH 8 in native cell lysate using no PLpro addition as negative control (cleavage events n = 59). Specificity profiles for P5-P5’ were determined as described in materials and methods. Positional occurrence of amino acids are shown as enrichment over expected frequency with natural amino acid abundance derived from Uniprot *homo sapiens* reference proteome UP000005640. Only enrichments >1.5-fold are displayed.

Four proteins were cleaved after glycine and one of them, heterogeneous nuclear ribonucleoprotein A2/B1 (HNRNPA2B1; UniProt Q9UKV8), was cleaved at a di-glycine motif (GGRGG216|YGGGG), closely resembling the canonical LRGG motif [12]. HNRNPA2B1 also contains basic-site cleavage at FGGSR_273_|NMGGP. Additionally, HNRNPA2B1 is reported to be cleaved by PLpro at the glycine-rich area of NQGG_281_|GYGG by Luo et al. (2023) [8]. An overview of all 59 significantly cleaved sequences is provided in Supplemental Table S3.

## 4. Conclusions

This study uncovers a previously unrecognised dibasic cleavage preference of the SARS-CoV-2 papain-like protease (PLpro) and shows that this signature is conserved across host proteomes and between the SARS-CoV-2 and SARS-CoV-1 enzymes. Using a PICS workflow with stringent TMT quantification, we defined a reproducible motif characterised by dominant arginine in P1, frequent arginine or lysine in P1′, and lysine as a secondary residue in P1, irrespective of the organismal background or enzyme source. The presence of the same motif in two related coronavirus proteases suggests evolutionary pressure to retain host-protein targeting beyond viral polyprotein processing.

Titration experiments revealed that PLpro’s turnover of basic sites is intrinsically low compared with trypsin, yet becomes more dominant once PLpro is supplied at higher enzyme-to-substrate ratios, indicating a genuine but secondary substrate class. Neutralising Asp-108, Glu-161 or Asp-164, three acidic residues flanking the catalytic triad, abolished prime-side arginine enrichment and inverted the P1 preference from arginine to lysine while modestly increasing overall activity. Although these coordinated shifts are consistent with a role for the acidic pocket in stabilizing arginine-rich junctions and reveal subsite cooperativity favoring cleavage between two basic residues, alternative explanations, such as subtle conformational changes affecting subsite architecture, cannot be excluded.

When the protease was applied to native HEK293 lysate, only 59 significant cut sites were detected, retaining arginine at P1/P1′ and lacking lysine. Prime-side sequences became Gly-rich, indicating a requirement for structural flexibility and suggesting that basic-site cleavage is restricted to accessible loops or termini in full-length proteins. Notably, heterogeneous nuclear ribonucleoprotein A2/B1 was cleaved at both di-glycine and dibasic motifs, illustrating dual binding modes.

Collectively, these results extend PLpro specificity beyond the canonical LXGG motif and establish a mechanistic link between an acidic surface patch and basic-residue recognition. The expanded cleavage-motif repertoire provides a framework for rational inhibitor design that targets both glycine- and arginine/lysine-binding modes and it offers a resource for predicting host pathways susceptible to coronavirus PLpro activity.

## CRediT authorship contribution statement

**Daniel Vogele**: Investigation, Formal analysis, Writing – Original Draft, Writing – Review & Editing, Visualization. **Klemens Fröhlich**: Methodology, Writing – Review & Editing. **Oğuz Bolgi**: Resources. **Ruth Geiss-Friedlander**: Resources, Funding acquisition. **Oliver Schilling**: Conceptualization, Writing – Review & Editing, Supervision, Funding acquisition.

## Data availability

The raw data used for peptide and protein identification and quantification, as well as the analysis output files and R scripts utilized in this study, are available via the Massive database (MassIVE MSV000097809) and the following FTP download link: ftp://massive-ftp.ucsd.edu/v09/MSV000097809/

## Acknowlegments

Oliver Schilling acknowledges funding by the Deutsche Forschungsgemeinschaft (DFG, projects 446058856, 466359513, 444936968, 405351425, 431336276, 431984000 (SFB 1453 “NephGen”), 441891347 (SFB 1479 “OncoEscape”), 423813989 (GRK 2606 “ProtPath”), 322977937 (GRK 2344 “MeInBio”)) 507957722, the ERA PerMed program (BMBF, 01KU1916, 01KU1915A), the German Consortium for Translational Cancer Research (project Impro-Rec), the MatrixCode research group, FRIAS, Freiburg, the investBW program BW1_1198/03, the ERA TransCan program (projects 01KT2201,”PREDICO”, 01KT2333 „ICC-STRAT”), the BMBF KMUi program (project 13GW0603E, project ESTHER), and the BMBF Cluster4Future program (nanodiag). Ruth Geiss-Friedlander acknowledges funding by the Deutsche Forschungsgemeinschaft GRK 2606 “ProtPath” project ID 423813989.

## Declaration of generative AI and AI-assisted technologies in the writing process

During the preparation of this work, the authors used ChatGPT for grammar checking and phrase refinement. After using this tool/service, the authors reviewed and edited the content as needed and take full responsibility for the content of the publication.

