## Supplemental Figures for "Specificity profiling of SARS-CoV-2 PLpro using proteome-derived libraries of linear peptides suggests secondary preference for basic motifs"

```
*
 BAD AVG GOOD
*
SARS-CoV-2      : 100
SARS-CoV-1      : 100
cons            : 100

SARS-CoV-2      EVRTIKVFTTVDNINLHTQVVDMSMTYGQQFGPTYLDGADVTKIKPHNSHEGKTFYVLPNDDTLRVEAF
SARS-CoV-1      EVKTIKVFTTVDNTNLHTQLVDMSMTYGQQFGPTYLDGADVTKIKPHVNHEGKTFFVLPSDDTLRSEAF

cons            **:********** *****:*************************** .******:***.***** ***


SARS-CoV-2      EYYHTTDPSFLGRYMSALNHTKKWKYPQVNGLTSIKWADNNCYLATALLTLQQIELKFNPPALQDAYYR
SARS-CoV-1      EYYHTLDESFLGRYMSALNHTKKWKFPQVGGLTSIKWADNNCYLSSVLLALQQLEVKFNAPALQEAYYR

cons            ***** * *****************:***.**************::.**:***:*:***.****:****


SARS-CoV-2      ARAGEAANFCALILAYCNKTVGELGDVRETMSYLFQHANLDSCKRVLNVVCKTCGQQQTTLKGVEAVMY
SARS-CoV-1      ARAGDAANFCALILAYSNKTVGELGDVRETMTHLLQHANLESAKRVLNVVCKHCGQKTTTLTGVEAVMY

cons            ****:***********.**************::*:*****:*.********* ***: ***.*******


SARS-CoV-2      MGTLSYEQFKKGVQIPCTCGKQATKYLVQQESPFVMMSAPPAQYELKHGTFTCASEYTGNYQCGHYKHI
SARS-CoV-1      MGTLSYDNLKTGVSIPCVCGRDATQYLVQQESSFVMMSAPPAEYKLQQGTFLCANEYTGNYQCGHYTHI

cons            ******:::*.**.***.**::**:*******.*********:*:*::*** **.***********.**


SARS-CoV-2      TSKETLYCIDGALLTKSSEYKGPITDVFYKENSYTTTIKPVTYK
SARS-CoV-1      TAKETLYRIDGAHLTKMSEYKGPVTDVFYKETSYTTTIKPVSYK

cons            *:***** **** *** ******:*******.*********:**
```

**Supplemental figure 1: T-Coffee multiple sequence alignment results of SARS-CoV-1 and SARS-CoV-2 PLpro.**
T-Coffee sequence alignment of SARS-CoV-2 PLpro and SARS-CoV-1 PLpro, with red highlighting indicating high alignment confidence. Asterisks (*) in the “cons” row denote identical residues, emphasizing the strong conservation between the two proteases (alignment score: 100).

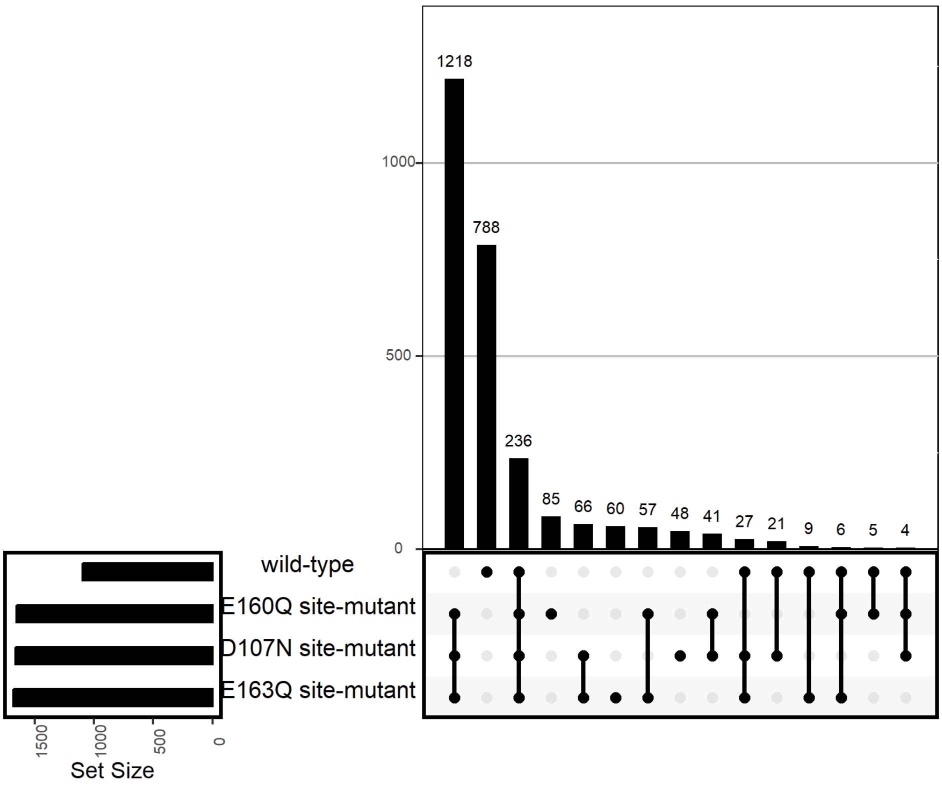


**Supplemental figure 2: Subset plot of SARS-CoV-2 PLpro wild-type and site mutant cleavage sequences.**
Set size illustrating the number of peptides with significantly increased cleavage-site abundance in SARS-CoV-2 PLpro wild-type and the D107N, E160Q and E163Q site-directed mutants. The top bar chart displays the size of each intersection (shared or unique peptide sets), and the matrix below indicates which conditions each peptide set is present in.

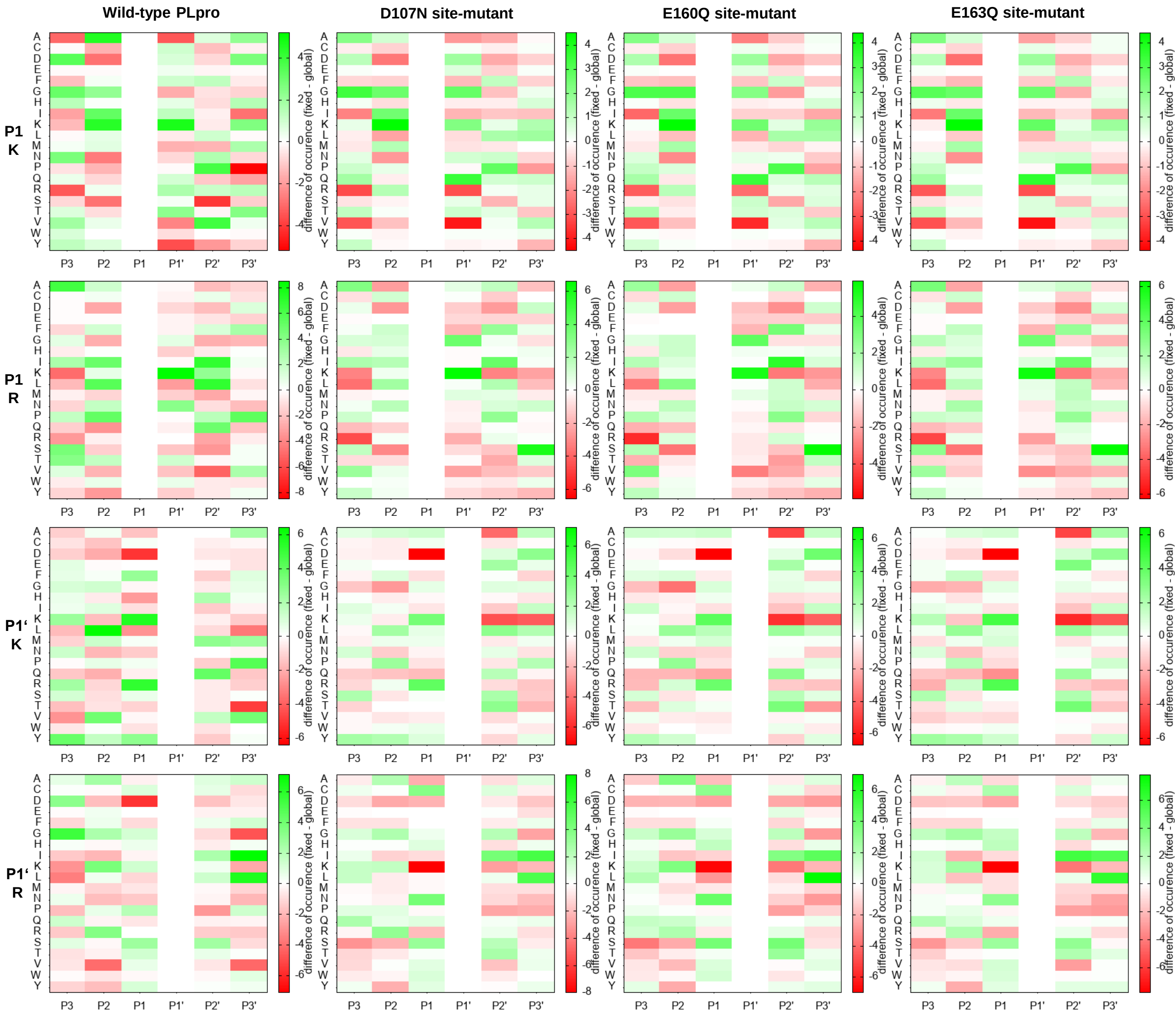


**Supplemental figure 3: Probing for putative sub-site cooperativity for SARS-CoV-2 PLpro and site mutant cleavage sequences with basic residues in P1 and P1’.**

Each subpanel depicts subsite cooperativity analysis for wild-type (WT) SARS-CoV-2 PLpro and site-mutants in cleavage sequences containing basic residues (K or R) in P1 and P1′. Rows represent amino acids, columns indicate positions (P3 to P1, P1′ to P3′), and the color scale indicates differences from a global reference frequency (red: depletion, green: enrichment).
